# Binding Mode of SARS-CoV2 Fusion Peptide to Human Cellular Membrane

**DOI:** 10.1101/2020.10.27.357350

**Authors:** Defne Gorgun, Muyun Lihan, Karan Kapoor, Emad Tajkhorshid

## Abstract

Infection of human cells by the SARS-CoV2 relies on its binding to a specific receptor and subsequent fusion of the viral and host cell membranes. The fusion peptide (FP), a short peptide segment in the spike protein, plays a central role in the initial penetration of the virus into the host cell membrane, followed by the fusion of the two membranes. Here, we use an array of molecular dynamics (MD) simulations taking advantage of the Highly Mobile Membrane Mimetic (HMMM) model, to investigate the interaction of the SARS-CoV2 FP with a lipid bilayer representing mammalian cellular membranes at an atomic level, and to characterize the membrane-bound form of the peptide. Six independent systems were generated by changing the initial positioning and orientation of the FP with respect to the membrane, and each system was simulated in five independent replicas, each for 300 ns. In 73% of the simulations, the FP reaches a stable, membrane-bound configuration where the peptide deeply penetrated into the membrane. Clustering of the results reveals three major membrane binding modes (binding modes 1-3) where binding mode 1 populates over half of the data points. Taking into account the sequence conservation among the viral FPs and the results of mutagenesis studies establishing the role of specific residues in the helical portion of the FP in membrane association, the significant depth of penetration of the whole peptide, and the dense population of the respective cluster, we propose that the most deeply inserted membrane-bound form (binding mode 1) represents more closely the biologically relevant form. Analysis of FP-lipid interactions shows the involvement of specific residues, previously described as the “fusion active core residues”, in membrane binding. Taken together, the results shed light on a key step involved in SARS-CoV2 infection with potential implications in designing novel inhibitors.

**Significance:** A key step in cellular infection by the SARS-CoV2 virus is its attachment to and penetration into the plasma membrane of human cells. These processes hinge upon the membrane interaction of the viral fusion peptide, a segment exposed by the spike protein upon its conformational changes after encountering the host cell. In this study, using molecular dynamics simulations, we describe how the fusion peptide from the SARS-CoV2 virus binds human cellular membranes and characterize, at an atomic level, lipid-protein interactions important for the stability of its membrane-bound state.

## Introduction

Coronavirus disease 2019 (COVID-19) emerged in late 2019 as a significant threat to human health. It became a global pandemic by March 2020,^1,2^ and it continues to claim lives and to significantly impact all aspects of people’s lives around the globe. COVID-19 is caused by severe acute respiratory syndrome coronavirus 2 (SARS-CoV2), a positive-strand RNA virus that causes severe respiratory complications, among other symptoms, in humans. ^3^ SARS-CoV2 recognizes and infects human cells that express a cell-surface receptor termed angiotensin-converting enzyme 2 (ACE2), ^4^ which is specifically recognized by the viral spike glycoprotein (S-protein). Binding of the two proteins is a prerequisite for the fusion of the viral and cellular membranes,^5^ one of the first and required steps in viral infection facilitating the release of the viral genome into the infected cell.^6–9^

Binding of the virus to the ACE2 receptor on the host cell is mediated by the S1 domain in the viral S-protein. The next key step, namely, virus-host membrane fusion is mediated by the S2 domain.^10^ S2 consists of multiple proteolytic cleavage sites, including one at the boundary of S1/S2 and one at the S2’ site, which are cleaved as part of the fusion process.^11–13^ Cleavages at these sites, downstream to the two heptad repeat regions (HR1 and HR2), induce the dissociation of the S1 subunits from the S-protein, followed by a series of conformational changes that trigger membrane fusion between the host cell membrane and the viral envelope. ^14,15^ The remaining S2 trimer, a post-fusion structural motif, is shared among all the class I viral fusion proteins. ^16,17^ A critical part of any viral fusion protein in the coronavirus family is the relatively hydrophobic fusion peptide (FP), a segment in the S2 domain which is responsible for directly interacting with and inserting into the host cell membrane, thereby initiating the fusion process.^8,18,19^

Viral FPs share several characteristics that help locating their position within their parent protein (the S-protien in SARS-CoV2, for example). The sequences of the FPs are highly conserved within each family of viruses (but not between families), the frequency of glycines and alanines in the sequence is relatively high, and their cleavage sites are often flanked by bulky hydrophobic residues as well as hydrophilic residues.^20,21^ Some FPs have a central kink produced by a proline and a helix-turn-helix structure, as observed in influenza virus hemagglutinin (HA). In such cases, proteolytic cleavage occurs directly at the N-terminus to the FP, and the peptides are thus called external or N-terminal FPs. In other cases, the proteolytic cleavage site resides upstream from the FP, which is relatively longer (25-30 amino acids) and contains a prolonged *α*-helix in the fusion-active state, e.g., in the cases of Ebola virus or avian leukosis sarcoma virus.^8,20,21^ Such FPs are referred to as internal FPs, which also is the case for SARS-CoV2.

To this date, three main FP regions have been proposed for SARS-CoV S-proteins,^22,23^ which are all located between the N-terminus and the HR1 region of the S2 domain: (1) at the N-terminus of HR1, (2) near the S1/S2 cleavage site, and, (3) at the C-terminus of the cleavage site S2’.^13^ Based on the criteria stated above as well as other experimental evidence, most recent data suggest that immediately downstream the S2’ cleavage site of the SARS-CoV2 is the leading segment involved in the fusion process. ^11,24–27^ Mutagenesis experiments showed the significance of the FP in this part of the protein, specifically terming a region the fusion-active core. ^21^

Cholesterol (CHL) is a major component of the human cellular membranes. ^28–31^ A recent fluorescence spectroscopy study has shown that CHL plays an important role in modulating the binding affinity and organization of the SARS-CoV FP in the membranes.^32^ Therefore, taking into account the natural lipid composition of a mammalian cell in simulation studies such as the present one is important.

Characterizing how the FP binds to the membrane and how it interacts with specific lipids has been challenging experimentally. Computational methods, particularly molecular dynamics (MD) simulation, offer an alternative strategy to capture the membrane binding process of the FP and to probe its binding mode to the cellular membrane. One of the major challenges in simulating such processes lies in sufficient sampling of possible FP membrane-binding poses. Due to the slow dynamics of membrane lipids, they are often insufficiently sampled on the timescales which atomistic MD simulations currently can access, causing the membrane binding and insertion of proteins to be biased by the initial lipid distribution and protein placement. In this context, an alternative membrane model, termed the highly mobile membrane-mimetic (HMMM) model, has been developed to enhance lipid diffusion without compromising the atomistic description of lipid head groups. ^33–36^ The HMMM model is based on the combination of a biphasic solvent system^33^ with short-tailed lipids at the interface.^34^ Owing to its substantially enhanced lipid mobility the model has proven extremely efficient in describing mixed lipid bilayers, reproducibly capturing spontaneous (unbiased) membrane binding and insertion of a wide spectrum of peripheral proteins,^34,37–48^ and collecting significantly improved sampling of lipid–protein interactions. Of particular interest to the present study, this method has also been successfully used to capture spontaneous membrane association of the influenza virus hemagglutinin FP.^49^

In this study, we perform an extensive set of HMMM simulations, to investigate membrane binding of the SARS-CoV2 FP, starting from several initial, different positioning of the peptide with respect to the membrane to further improve sampling. The results provide a detailed mechanistic picture of the initial step in the fusion process, focusing on the biophysical aspects of the virus–lipid bilayer interactions during this process. Characterizing the mechanism of the fusion-driving, FP-host membrane interactions is key to our understanding of critical steps involved in viral infection, and might pave the way for development of novel therapeutic interventions and strategies against the virus.

## MATERIALS AND METHODS

### Modeling of the SARS-CoV2 fusion peptide (FP)

As the first step for modeling the SARS-CoV2 FP, multiple sequence alignment was carried out for the S-proteins from different human coronaviruses (HCoVs). There are a total of seven known HCoVs: 229E and NL63 from the alpha subfamily, and OC43, HKU1, Middle East respiratory syndrome (MERS), severe acute respiratory syndrome (SARS) and SARS-CoV2, from the beta subfamily. Out of these, S-protein structures are available for HKU1, MERS, SARS, as well as SARS-CoV2. Additionally, bat coronavirus RaTG13, a closely related homolog of SARS-CoV2, was also included in the sequence alignment. Multiple sequence alignment for the above eight sequences was carried out using the MAFFT program with the L-INS-i method^50^ and visualized using Jalview^51^ (Fig. S1). The sequence alignment highlighted the high degree of residue conservation among the different coronaviruses, especially in the FP region, and informed the FP structural modeling as described below.

The available cryoEM structure of the SARS-CoV2 S-protein at the time (PDB: 6VSB)^52^ contained only a partial structure (12 residues) of the FP, making it unsuitable for construction the initial SARS-CoV2 FP model (Fig. 1A). Since the S2 domain of the S-protein containing the FP is well-conserved among the SARS coronaviruses, we used the S-protein from SARS-CoV as a template for modeling the FP of the SARS-CoV2. Accordingly, the cryoEM structure of the S-protein from SARS-CoV (PDB: 5XLR)^53^ containing the FP structure was used as a template for constructing the initial SARS-CoV2 FP model. Our SARS-CoV2 FP model also contained the loop connecting the FP to the neighboring proximal region, suggested to be important in the fusion process,^24–26^ but not modeled fully in the current study due to lack of a suitable structural template at the time. The only two different residues between the sequences of SARS-CoV and SARS-CoV2 FPs (I834/M816 and D839/E821 in Fig. S1) were mutated to the SARS-CoV2 residues (Fig. 1B).

**Figure 1:**
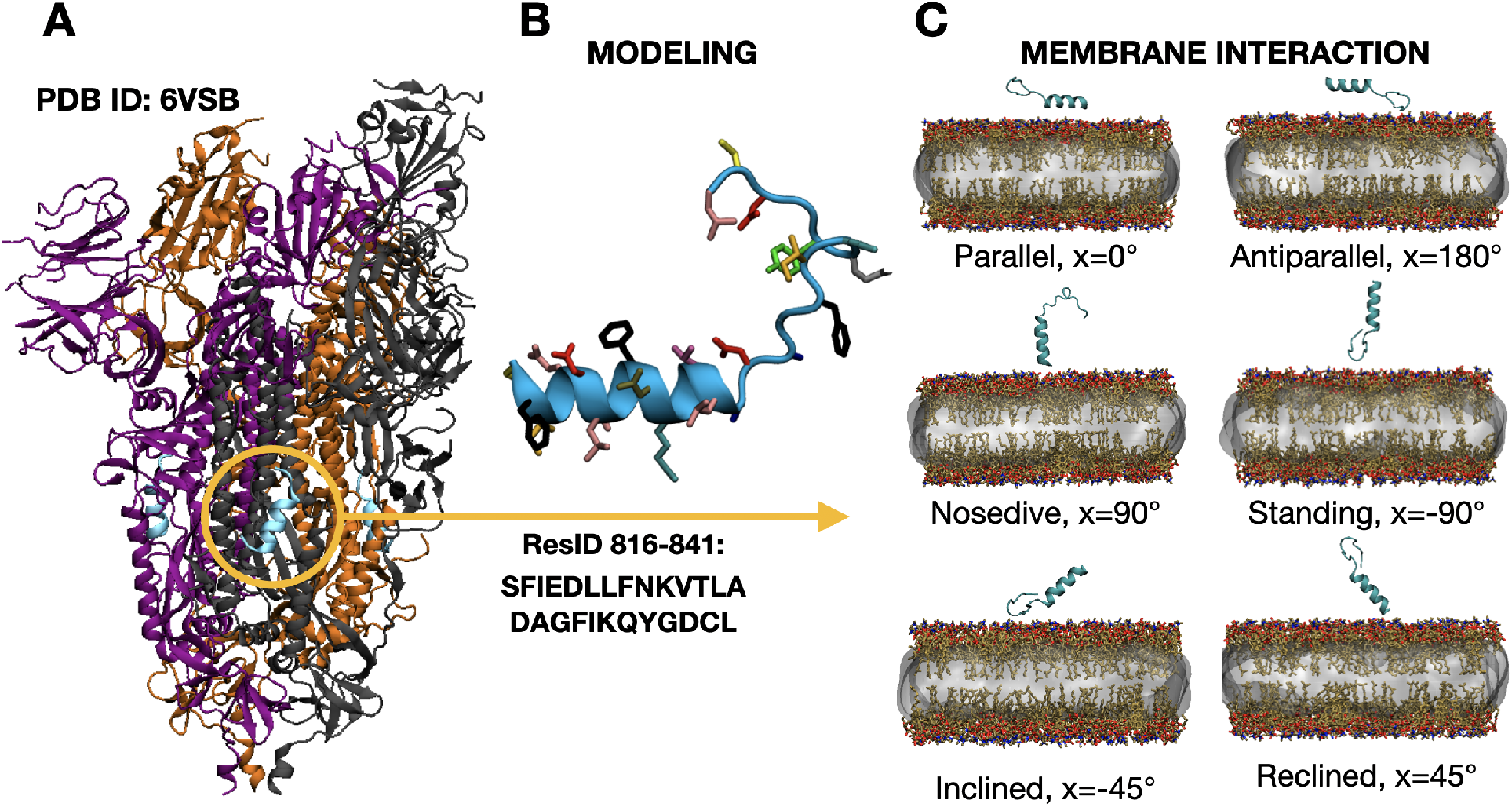
Modeling and design of membrane binding simulations of the SARS-CoV2 FP. A) Cryo-EM structure of pre-fusion trimeric SARS-CoV2 S-protein with the highlighted FP (blue). Each monomer is drawn in a different color (grey, purple, and orange). B) The SARS-CoV2 FP. The missing residues of the SARS-CoV2 FP structure were modeled using SARS-CoV FP (PDB 5XLR), with the mutations of I834/M816 and D839/E821 from SARS-CoV to SARS-CoV2 (Fig. S1). Residues 817-828 are in an alpha helical conformation and residues 829-841 form a loop. C) HMMM membrane binding simulation setup. After simulating the modeled FP in solution for 20 ns, the equilibrated peptide was placed above the HMMM lipid bilayers in several different orientations. The membrane lipid composition is PC/PE/CHL (60/6/34) mol% which are major lipids composing outer leaflet of the human plasma membrane (carbon as tan, oxygen as red, phosphorus as blue, and nitrogen as orange). We use six different initial orientations rotating the peptide around its x axis with respect the parallel orientation: parallel (P, x=0°), antiparallel (A, x=180°), nosedive (N, x=90°), standing (S, x= −90°), inclined (I, x= −45°), and reclined (R, x=45°)), for the FP placement above the membrane to randomize the initial orientation. Lipid bilayers were randomly and individually built for each replica. Each orientation was simulated in five independent replicas, with the total replica number of 30 for membrane-binding simulations.

The initial FP model was then solvated and ionized with 0.15 M NaCl using SOLVATE and AUTOIONIZE plugins of VMD, respectively). ^54^ Energy minimization was carried out using the steepest descent method for 100 timesteps followed by an equilibrium MD of the FP in solution for 20 ns in an NPT ensemble, in order to obtain a fully relaxed initial SARS-CoV2 FP in an aqueous environment. The resulting equilibrated FP was used in the subsequent membrane-binding simulations.

### Membrane preparation

To obtain membrane-bound models of the SARS-CoV2 FP, we performed multiple independent simulations employing the HMMM model (Fig. 1C) ,^33,34,36^ in which the diffusion of lipids is enhanced allowing for better sampling and convergence of FP–lipid interactions. As shown in previous studies, compared to conventional (full) membrane simulations, physical characteristics of membrane binding remain conserved for peripheral proteins but the binding timescale is reduced by an order of magnitude with the use of HMMM.^38,41,55^ The HMMM model has been extensively used in a broad spectrum of systems providing mechanistic details of lipid-protein interactions^37–41,43–49,56–65^ and can accurately reproduce the energetics associated with partitioning of amino acids within the head groups and in the most peripheral section of the leaflets.^35^ We use this technique to facilitate binding and insertion of the FP into the membrane, which would allow us to focus our computational effort on running many independent membrane-binding simulations.

In the present study, symmetric full membranes were first constructed using CHARMM-GUI,^66^ with a lipid composition (POPC/POPE/CHL: 60/6/34 mol%) that included major lipids in the outer leaflet of the human plasma membrane,^28–31^ except for sphingomyelin, which has not been sufficiently tested in a cholesterol-containing HMMM model. The initial, full membranes generated with CHARMM-GUI were converted to HMMM membranes by removing the atoms after the fifth carbon in the phospholipid acyl chains, resulting in short-tailed PC and PE lipids, while keeping the cholesterol molecules intact. To mimic the membrane core, a previously developed *in silico* solvent, termed SCSE (including two carbon-like interaction centers),^36^ was used to match the number of heavy atoms removed from the lipid tails in the previous step. The resulting HMMM membranes contained 2,178 SCSE molecules^36^ and 150 lipids in each leaflet.

In order to further expand the sampling of the phase space, we varied the initial placement and orientation of the FP, i.e., six rotated orientations (P, A, N, S, I, R) (Fig. 1C), to minimize the initial bias. The systems were solvated and ionized using the SOLVATE and AUTOIONIZE plugins in VMD, ^54^ with 0.15 M NaCl resulting in total system sizes of approximately 70,000-90,000 atoms and box sizes of 103×103×90 to 103×103×110 Å^3^. Multiple replicas were simulated for each FP orientation, as the diffusion and mixing of lipids and the process of membrane binding and insertion of the FP can be slow even when using HMMM membranes, and more sampling would ensure the reliability of the obtained membrane-bound configurations. Five independent HMMM membranes with each specified FP orientation were generated using a Monte Carlo based lipid mixing protocol developed in our group to further enhance variation of initial membrane configurations for each FP orientation.

### Membrane-binding simulations

The systems were energy-minimized using steepest descent method for 10,000 steps and simulated for 10 ns with the C*α* atoms of the peptide harmonically restrained (k = 0.5 kcal mol^−1^ Å^−2^), followed by a production run of 300 ns after removing the C*α* restraints. A harmonic restraint along the z axis, with a force constant of *k* = 0.05 kcal mol^−1^ Å^−2^ was applied to the C2, C26, and C36 atoms of the phospholipids, and to the O3 atom of cholesterol, to reproduce the atomic distributions of these lipids in a full lipid bilayer more closely, and to prevent the occasional escape of short-tailed lipids into the aqueous phase, which is expected for these surfactant-like molecules. To prevent SCSE molecules from diffusing out of the core of the membrane, we subjected them to a grid-based restraining potential, applied using the gridForce^67^ feature of NAMD. ^68,69^ Five replicas, each with an independently generated HMMM membrane (different lipid mixings) and a starting orientation of the FP, were simulated, resulting in a total of 30 independent membrane-binding simulations for a cumulative time of 9 *μs*. The initial and final structures for all the simulations, as well as representative membrane-bound structures, topology/parameter files, and configuration files for the runs are all made available in Open Science Frame-work (OSF), https://osf.io/qpfds/?view_only=810f1d0cad3140c690bb0d4fa50e0afe. Trajectory files were prohibitively large to be deposited, but will be made available upon request.

### Simulation protocol

The systems were simulated using an NPT ensemble with a constant x/y ratio, using a target pressure and temperature of 1.0 atm and 310 K, respectively, and a time step of 2 fs. Although scaling up the area combined with a constant-area protocol is commonly used for HMMM simulations to allow faster membrane binding,^33,34,36^ given that the FP is considerably small, we did not expect that our NPT setup would significantly hamper the process of membrane binding. Simulations were performed using NAMD^68,69^ with the CHARMM36^70^ force field parameters for lipids and proteins. The TIP3P model was used for water.^71^ The Nosé-Hoover Langevin piston method^72,73^ was utilized to maintain the constant pressure, and constant temperature was maintained by Langevin dynamics with a damping coefficient *γ* of 0.5 ps ^−1^ applied to all atoms. Nonbonded interactions were cut off after 12 Å starting at a switching distance of 10 Å. The particle mesh Ewald (PME) method^74^ was used for long-range electrostatic calculations with a grid density greater than 1 Å^−3^.

### Analysis

We define stable membrane binding with the criteria described below (see Fig. 3). First, a contact between the FP and the membrane lipids is defined for any heavy atom in the FP that is within 3.5 Å of any lipid heavy atom. Any contiguous segment of the simulation trajectory with a length of at least 30 ns during which at least one contact between the lipids and the FP existed was considered stable binding (marked with a blue background in Fig.3).

To characterize the binding orientation of the FP with respect to the membrane, the first principal axis (PA) of the FP helical segment (residues 816-823) was calculated (Fig. S2). The angle between the first PA and the membrane normal, *θ*_PA_, was used to represent the tilting of the FP. We then identify two additional angles to describe the orientation of two phenylalanine residues on the FP helical segment with respect to the membrane: *θ*_F817_ and *θ*_F823_, which are defined by the angle between the membrane normal and the perpendicular component (to the PA) of F817 and F823 C*α*-C*β* vector, respectively. Combining *θ*_PA_ with these two additional angles, which take into account the rotational degrees of freedom of the FP, we obtain a full description of the PF binding orientation to the membrane.

For all the frames identified as membrane-bound (see above for definition), a vector composed of the *z*-distances between the centers of masses (COMs) of individual FP side chains and the lipid phosphate plane was calculated. A dissimilarity matrix was then constructed using the euclidean pairwise distance of this vector representation for each frame. The *k*-medoids clustering algorithm^75^ was then performed to classify the membrane-bound poses of the FP, resulting in three major membrane binding clusters. The approximate center, i.e., the medoid, was used as the representative structure for each cluster, that is, for each identified FP membrane binding mode. To better visualize the the clustering results and the distribution of membrane-bound poses, principal component analysis (PCA) was performed to reduce the dimensionality of the original data set. In PCA, the covariance matrix of the *z*-distances between the COM of individual residues and the phosphate plane was computed and diagonalized. The resulting eigenvectors, i.e., the principal components (PCs), represent the coordinates that maximize the variance of the projected data. The first two PCs were selected to project the original *z*-distance data of membrane-bound frames along with the clustering result onto the reduced dimension. The PCA was performed using the scikit-learn package.^76^

## RESULTS AND DISCUSSION

The SARS-CoV2 attachment to and its penetration into the human plasma membrane are key steps in its viral infection. These steps rely on the interaction of the viral FP with the host cellular membrane. This aspect is the central phenomenon that we study here using atomistic simulations, in order to characterize the binding pose and the conformation of the SARS-CoV2 FP when bound to the membrane. In the following sections we first describe the results of our membrane binding simulations in terms of the overall binding of the FP using some coarse parameters. Then, we use the depth of insertion of individual FP residues (COMs) in the membrane in a clustering analysis. We then examine the clustering results in a reduced dimension, offered by PCA, to better classify the different microstates that arise during the membrane-binding simulations and discuss the biological relevance of the resulting bound states.

### Spontaneous Membrane Binding and Insertion of SARS-CoV2 FP

In order to characterize the membrane binding mode of the SARS-CoV2 FP to human cellular membranes, we performed 30 independent membrane-binding simulations using HMMM lipid bilayers. Spontaneous diffusion and membrane binding of the FP in each simulation replica can be monitored by tracking the position of the center of mass (COM) of the peptide with respect to the membrane, as a coarse metric. The *z* component of the COM was tracked with respect to the phosphate layer (the average *z* position of all the phosphate groups) of each leaflet (blue and red lines, respectively, in Fig. 2). Due to the applied periodic boundary conditions, and the free diffusion of the FP in the solution, the FP was able to diffuse towards either the upper or the lower leaflet of the membrane (Fig. 2), both containing lipid compositions representing the outer leaflet of human plasma membrane.^28–31,77^ Among the 30 performed membrane-binding simulations, instances of both stable (majority) and transient binding events, as well as cases with no membrane binding, were observed. Since the lipids in our simulated membranes are all neutral, we do not expect any major electrostatic forces driving the binding. Rather, the major driving forces between the two elements are hydrophobic effects, and the FP diffusion in the solution can make the peptide take a longer time to make the initial encounter with the membrane.

**Figure 2:**
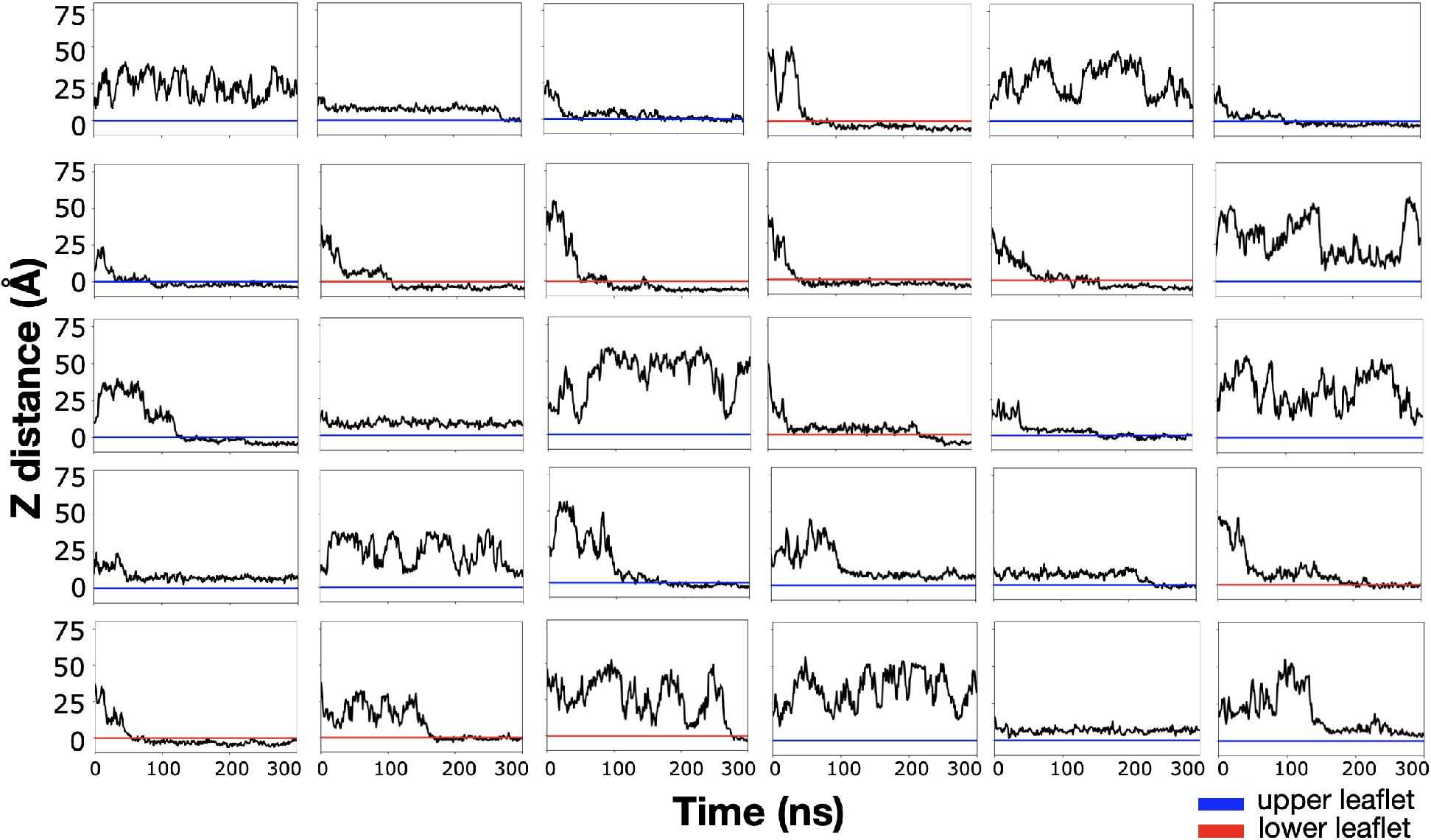
Spontaneous membrane binding of the SARS-CoV2 FP during the 30 independent membrane-binding simulations. The *z* component of the center of mass (COM) of the FP with respect to the average *z* position of the lipid phosphates in each leaflet, referred to as the phosphate layer. Replicas where the FP interacts with the lower leaflet are shown with a phosphate layer colored in red, and those binding to the upper leaflet in blue (due to periodicity binding to either leaflet is possible).

For further analysis of the membrane interactions, we selected only the portions of the trajectories where “stable membrane binding” or a “membrane-bound state” was defined (see Methods for details). Contacts between the FP and the lipid bilayer are shown in Fig. 3 where the blue segments of the graphs are considered stable binding (lipid contact for at least 30 ns). In 22 of the 30 independent membrane-binding simulations (~73%) membrane-bound configurations are observed (Fig. 3). For example, we observe stable membrane binding in parallel replicas P2, P3, P4, and P5, and in antiparallel replicas A1, A2, A3, and A5 (Fig. 3). In some simulations, nearly the entire length of the FP was observed to be engaged with membrane lipids (e.g., replicas P2, P5, A1, N1, I4, I5 and R1), while in other cases, only a specific part of the peptide makes contact with the membrane. Randomizing the initial placement of the peptides prevented biasing and allowed for better sampling the membrane interaction of the FP. In some simulations, the FP bound to the lipid bilayer as early as in 10 ns, while in the others it diffused and tumbled longer prior to interacting with the membrane, naturally allowing the peptide to further decouple itself from the effect of initial placement. For example, the initially reclined peptide in simulation replica R5 results in a membrane-bound configuration in which the FP interacts with the lipids through its loop part, or initially standing-placed peptides in S1 and S2 simulations end up being buried in the bilayer via their helical segment (Fig. 3).

**Figure 3:**
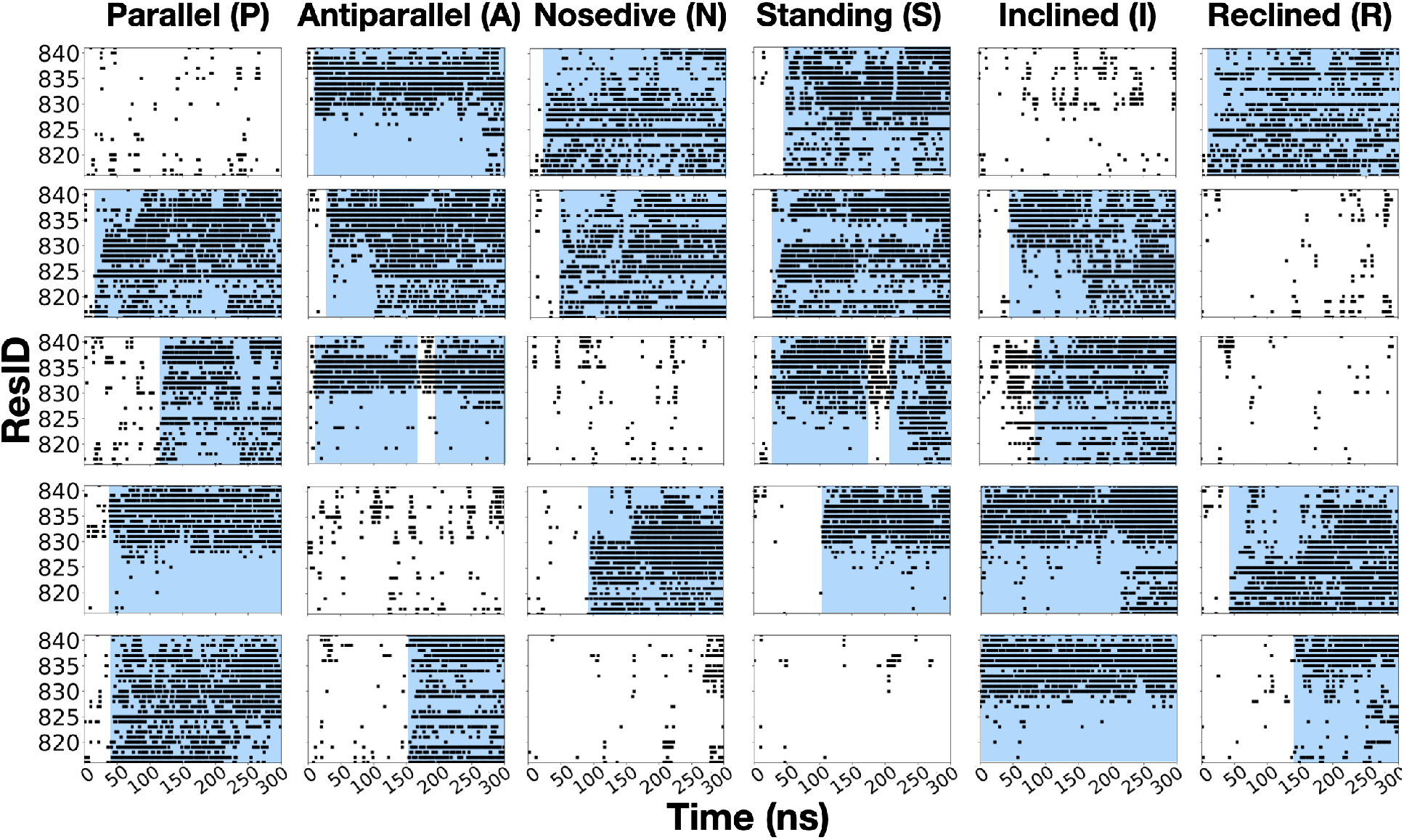
Identifying stable membrane binding using FP-lipid contacts. The plots show contacts between individual FP residues (y axis) and the lipids (heavy atom-distance of less than 3.5 Å) over time (x-axis) in a sampling step size of 1 ns. All contacts are labeled with black dots. The segments of the trajectories in which, for a minimum of 30 ns, at least one contact existed between the FP and lipids are marked with a blue background. We observe stable binding in 22 out of 30 replicas.

The overall structure of the FP plays a major role in its mode and depth of membrane binding. Therefore, we monitored the structural evolution of the FP throughout the simulations. The N-terminal *α*-helical segment (residues 816-823) remains largely unperturbed during the simulations, whereas the rest of the FP structure undergoes conformational changes. Within the membrane-bound configurations, the most common FP structure is still a hybrid *α*-helix/loop structure in which either the initial structure is mostly preserved or the *α*-helical segment is extended (Fig. S4B, Panels 2 and 5), which has been experimentally observed in many viral FPs when inserted into the membrane, for example for the case of Herpes and Hendra viruses. ^78–80^ In some replicas, we observed FPs evolving into hairpin-like structures (helix-hinge-helix), as shown, e.g., in Panels 1 and 2 of Fig. S4B, another structural arrangement also commonly reported for viral FPs. ^81,82^ In few cases, we also observed partial unfolding of the *α*-helical segment (Fig. S4B, Panel 4). From an energetic perspective, insertion of helical segments into the membrane might be less costly, since backbone amide groups in a helix can satisfy their hydrogen bonds internally, ^83^ which explains the preservation and even extension/formation of helical segments in most of our membrane-binding simulations. Since the secondary structure evolves in time, hereon for simplicity, the terms *α*-helical segment and loop segment will refer to the initial *α*-helix (residues 817-827) and loop (residues 828-841) parts of the FP.

Using the stable membrane-bound states defined above, we further examined the average position and lipid interaction of individual membrane bound FP residues (Fig 4). The average depth of insertion of individual residues reaches as deep as 10 Å below the phosphate layer, indicating interaction with the hydrophobic core of the membrane. From the pattern of residue–lipid interactions, we observe three major classes of membrane-bound states where either the whole peptide, the *α*-helical segment, or the loop part of the FP, interacts with the lipid bilayer, respectively. For example, in replicas P3, P5, and A5, the FP is almost completely buried into the membrane, in replicas P2, N1, and N2, the FP is bound to the membrane primarily through its *α*-helical segment, and in replicas P4, A1, A2, and A3, the loop is mostly engaged with the lipids. In order to better classify these binding modes in a reduced space, we next performed clustering of the obtained membrane-bound configurations.

**Figure 4:**
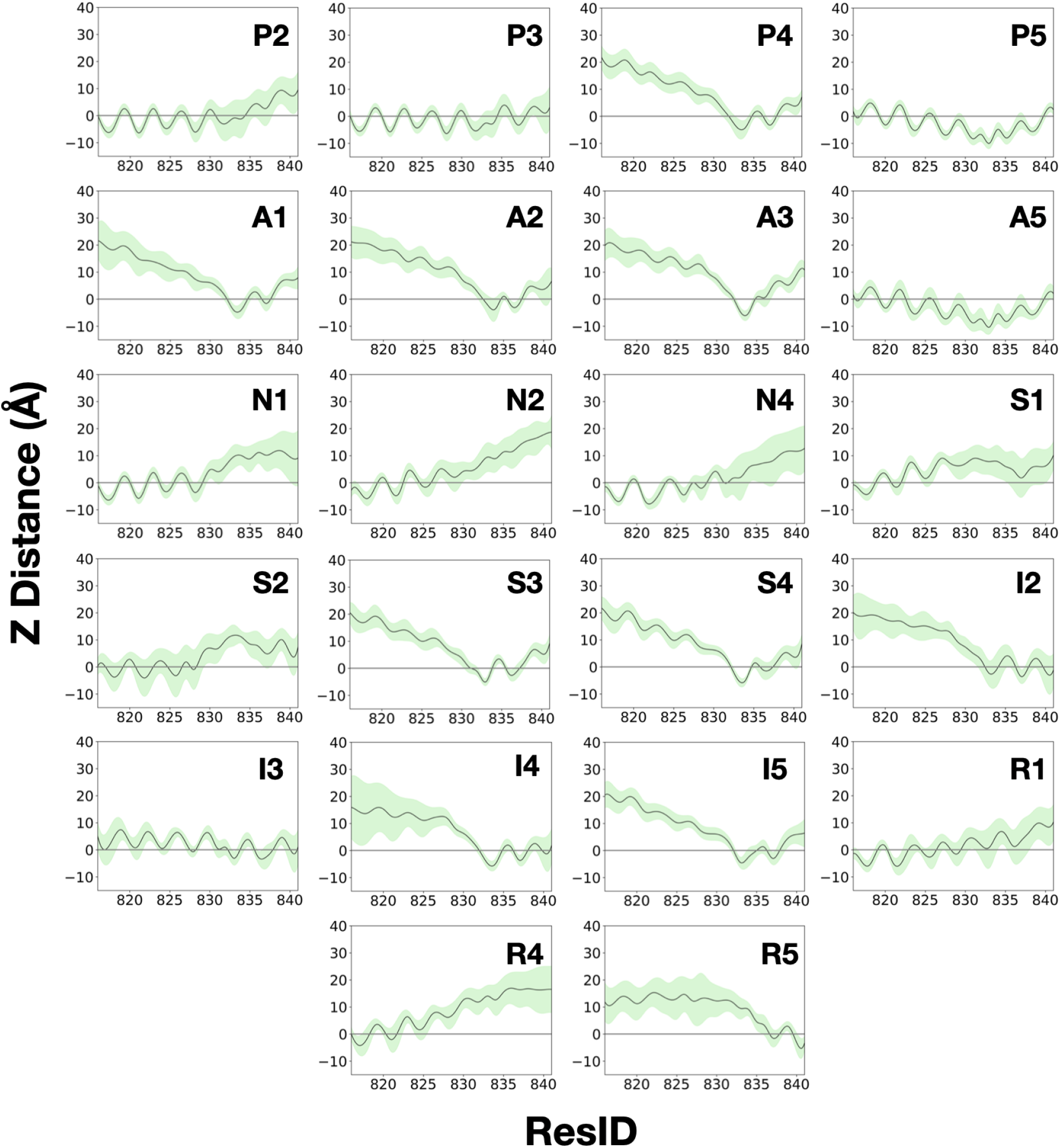
Insertion of FP residues into the membrane. Plots show the ensemble average of the side-chain COM *z* with respect to the phosphate plane of the bilayer in replicas with stable FP–membrane binding. Each data point is the average from the stably bound segments of each individual trajectory (blue-shaded areas in Fig. 3). Transparent green areas represent the standard deviation of the distance.

### Clustering and characterization of FP binding modes

To analyze the ensemble of FP membrane-bound configurations captured in the 22 HMMM simulations where stable membrane binding was observed, clustering was performed using the insertion depth of individual side chains with respect to the membrane (see Methods for details). Despite the large variance in membrane-bound poses among MD snapshots from different simulation replicas, three major clusters can be identified, which represent the three distinct binding patterns mentioned briefly above. The three binding modes (termed 1, 2, and 3) make up 50.4%, 11.2% and 38.4%, respectively, of the whole population of membrane-bound configurations observed in the simulations.

For better visualization, we performed PCA and reduced the dimension to the first two PCs, which together cover 82.4% of the variance (Fig. 5A). The nature of these two PCs, PC1 and PC2, was examined by their eigenvectors, which quantitatively evaluate the contribution of membrane insertion by each residue (Fig. S3). Based on the eigenvectors, PC1 and PC2 can be viewed as a measure of the membrane insertion of the helical segment (PC1) and that of the the C-terminal loop (PC2), with more negative values indicating deeper insertion. The clustering centers, i.e., the medoids, of the three binding modes are selected as representative binding poses (Fig. 5B, C, D). The three selected poses here represent the most sampled forms (clusters) of membrane-bound FP, and thus the energetically most favored states. The convergence of these three binding states can be observed from MD snapshots that are closest to the clustering centers while at the same time belonging to different replicas (Fig. S5). Binding mode 1, which accounts for over half of the data, corresponds to a pattern where the whole FP is deeply buried into the membrane (Fig. 5B). Binding modes 2 and 3 (Fig. 5C and D, respectively) represent membrane-bound configurations with either the helical segment or the C-terminal loop segment being the primary membrane-interacting region, respectively. We note that there are snapshots of membrane-bound FP poses located at the boundaries between the clusters or at the far edge of each cluster, which are much less sampled compared to the most representative region of the cluster. We believe these snapshots correspond to intermediate membrane-bound poses that are less biologically relevant.

**Figure 5:**
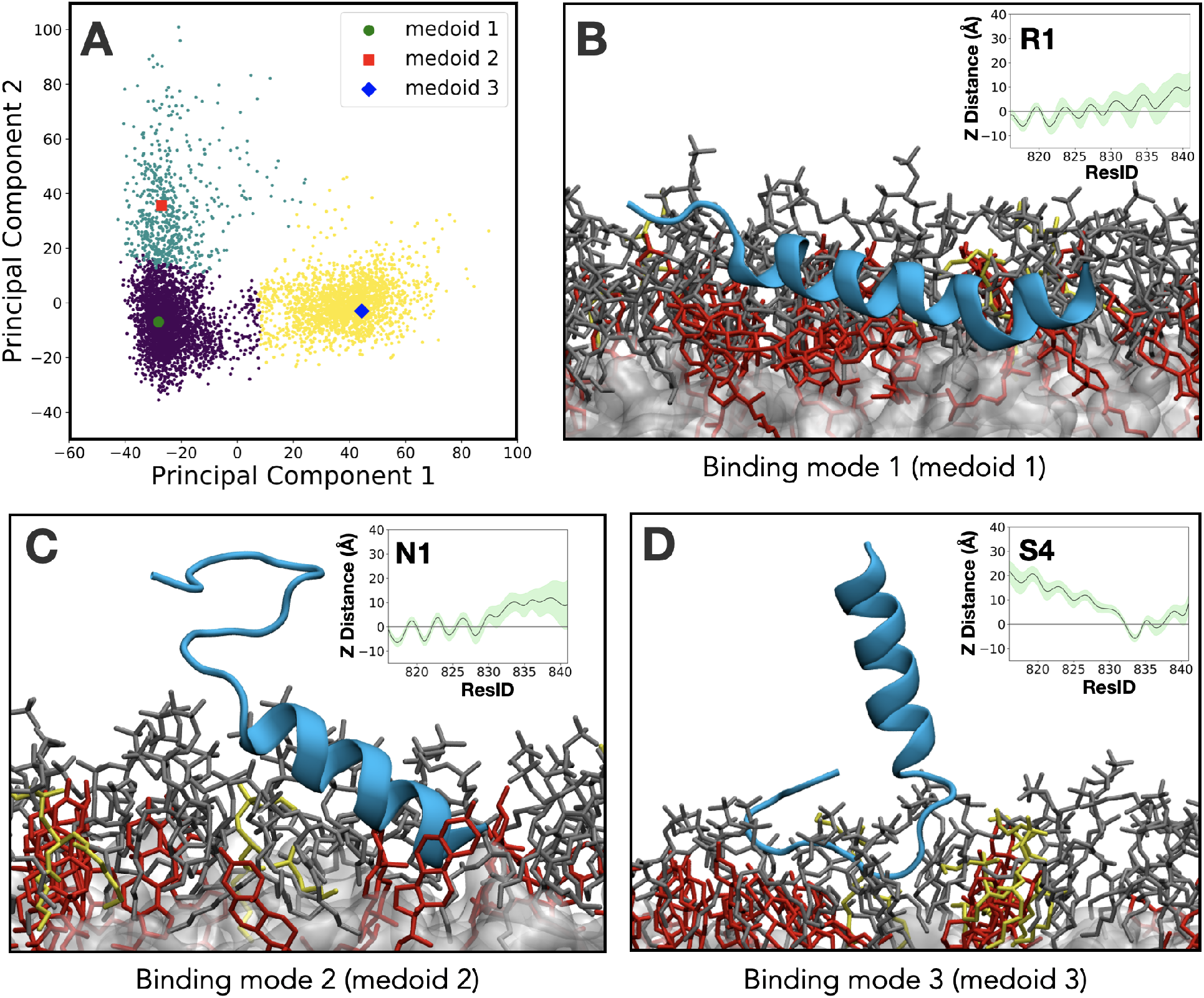
Clustering analysis of membrane-bound SARS-CoV2 FP and representative binding modes. A) Membrane-bound poses projected on PC1 and PC2. The binding modes/clusters 1, 2, and 3 are colored in purple, cyan and yellow with their clustering centers highlighted in green circle, red square and blue diamond, respectively. B) Representative binding mode 1: the binding configuration where the whole peptide is deeply buried into the lipid bilayer (at *t* = 230 ns in simulation R1). C) Representative binding mode 2: the configuration where the helical segment interacts with the bilayer in an oblique manner (*t* = 28 ns in simulation N1). D) Representative binding mode 3: the binding mode where the loop is inserted in the membrane (*t* = 123 ns in simulation S4). The insets in B, C, and D show the COM distance of each residue with respect to the phosphate layer, averaged over all the membrane-bound states in the trajectory.

### Orientation of the FP in its Membrane Bound Configuration

To further analyze the membrane binding configurations for SARS-CoV2 FP in our simulations, we calculated the orientations of the FP throughout the simulations (Fig. S2). We defined three angles, *θ*_PA_, *θ*_F817_, and *θ*_F823_, describing the FP helix orientation (see Methods for the definition of the angles) over the simulation trajectories (Fig. 6A) in replicas showing stably bound configurations. Although occasionally the FP can switch between binding modes 1, 2 and 3 within the same simulation replica, the analysis of these angles in different replicas will provide a better qualitative description of major binding modes observed in our simulations. For the replicas where binding mode 1 is the dominant binding configuration (e.g., P3, P5, A5, I3, R1), the average *θ*_PA_ angle is 90.4° (Fig. 6B), which corresponds to a parallel orientation with respect to the membrane. In replicas where binding mode 2 is dominant (N1, N2, N4, R4), *θ*_PA_ averages at 104.7° (fig. 6B), indicating an oblique configuration with the N-terminus facing the membrane core. Since the C-terminal segment is more flexible and the orientation angles are defined for the *α*-helical segment, it is more difficult to provide an equally clear description for binding mode 3 (e.g., P4, A1, A2, A3, S3, S4, I2, I4, I5). For this binding mode, in most of the replicas towards the end of the trajectory, *θ*_PA_ is less than 90° averaging at 51°. This means that the N-terminal of the helix is facing up, and as the angle decreases the helix becomes more orthogonal to the membrane. The *θ*_F817_ angle in binding modes 1 and 2 stabilized at average values of 134.6° and 137.2°, respectively, with a peak around 160°, where F817 is facing almost directly the membrane core (Fig. 6B). F823 is located on the opposite side of the helix with respect to the F817, therefore, in binding modes 1 and 2 this residue is mostly facing up and away from the membrane with average angles of 53.6° and 59.6°, respectively.

**Figure 6:**
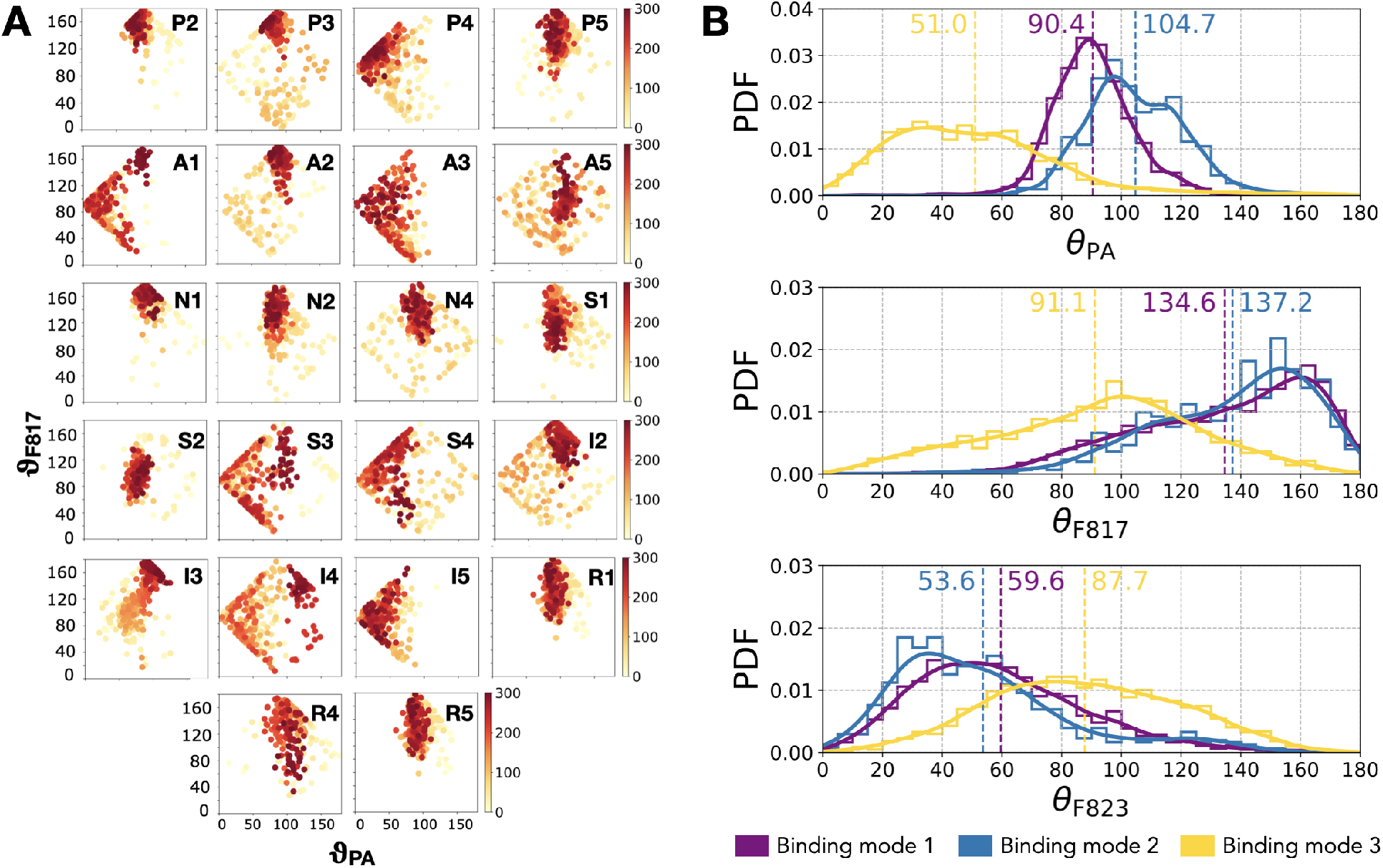
Orientation of the FP. A) Time evolution of two of the angles, *θ*_PA_ and *θ*_F817_ (Fig. S2), used to define the orientation of the FP with respect to the membrane. The color bars (from white to red) indicate the simulation time in ns. B) Probability density function (PDF) of FP’s three orientation angles (*θ*_PA_, *θ*_F817_, and *θ*_F823_) in the three major binding modes (clusters), colored in purple, cyan, and yellow, respectively. The PDFs (solid) are estimated by Gaussian kernel density estimation from the normalized histograms (stairstep). Average values are shown as dashed vertical lines and also labeled in the figure.

### Physiologically Relevant Membrane-Bound Configuration of SARS-CoV2 FP

In order to discuss our results within the a biology/physiology context, we compare our findings with experimental observations. We also assess the mechanistic relevance of the observed binding modes within the context of the whole S-protein by evaluating the way specific FP residues interact with the membrane.

In our simulations we observe three major membrane binding modes for SARS-CoV2 FP: binding mode 1, in which the peptide is nearly parallel to deeply buried in the membrane, binding mode 2 where the the helical segment of the FP is the primary site engaging with the membrane, and binding mode 3 where the C-terminal of the peptide is primarily interacting with the membrane (Fig. 5). Mutagenesis experiments have shown that highly conserved residues L821, L822, and F823, termed the “fusion active core”, play a major role in viral fusion of SARS-CoV .^21,84^ Given the high sequence similarity between SARS-CoV and SARS-CoV2 FPs, including L821/L822/F823 (Fig. S1), the fusion active core is likely to be also crucial for the binding and insertion of SARS-CoV2 FP. Consistent with these results, in binding modes 1 and 2, residues L821, L822, and F823 are deeply inserted into the hydrophobic core of the membrane, closely interacting with the membrane lipids (Figs. 5C and 7A). Additionally, electron spin resonance (ESR) spectroscopy experiments have shown that the highly conserved FP in SARS-CoV forms a “fusion platform”. In this predicted structure, the majority of the FP residues are deeply inserted into the membrane whereas C-terminal residues are free in the solution, ^24,25^ in a similar fashion to what we observe in binding mode 1 (Fig. 7A).

**Figure 7:**
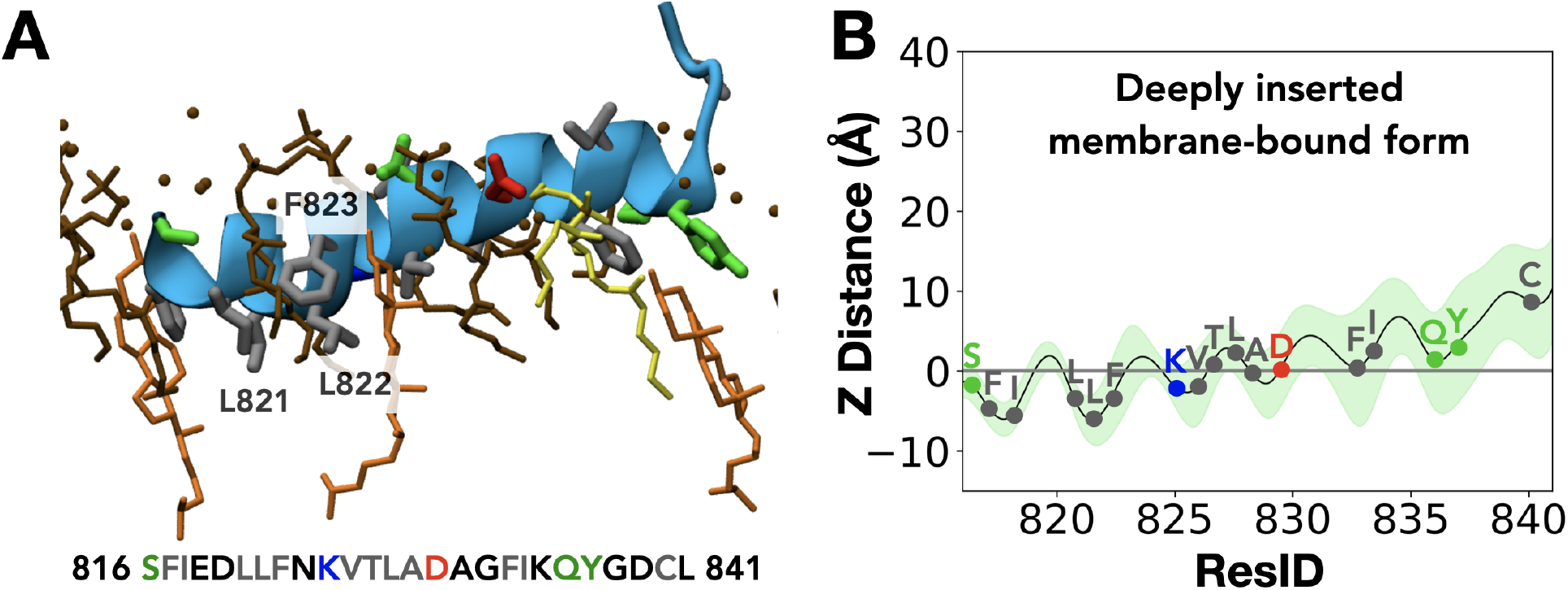
Deeply inserted membrane-bound FP representing the most physiologically relevant membrane-bound form. A) Deeply inserted membrane-bound form is shown in a detailed manner. Directly interacting lipids (heavy atoms closer than 3.5 Å) such as cholesterol (orange), PE (yellow), and PC (brown), as well as lipid-interacting side chains (hydrophobic: grey; acidic: red; basic: blue; and polar: green). Phosphorus atoms of phospholipids are shown as brown spheres. B) Average z-distance of each residue COM with respect to the phosphate bilayer is shown for the representative structure of binding mode 1, with the position of the residues directly involved in membrane interaction (heavy atom distance equal or closer than 3.5 Å marked, with coloring consistent with Panel A.

Evaluating the binding positions of the FP in the context of the post-fusion S-protein is another important component in identifying the most physiologically relevant binding form of the FP. The cleavage taking place prior to and aiding the conformational change of the S-protein at the the S2’ cleavage site makes S816 the N-terminus of the whole post-fusion S-protein. Therefore, FP in binding modes 1 and 2, which both enter the membrane with their N-terminus facing the membrane core at an oblique angle, are mechanistically viable and compatible with the nature of the predicted conformational changes in the S-protein. This is in sharp contrast to binding mode 3 where the C-terminal segment is buried in the membrane.

Furthermore, binding mode 1 is the most highly sampled binding mode in our simulations, accounting for more than half of the membrane-bound forms we observe in our entire data set. It is also the most deeply inserted binding form among the three major bound configurations. Given the established role of several hydrophobic residues in the helical segment of the FP in membrane interaction, the resemblance to the fusion platform, and its depth of penetration in the membrane, we strongly believe and propose that the binding mode 1 represents the physiologically relevant membrane-bound form of SARS-CoV2 FP, which we term here as the deeply inserted membrane-bound form.

The deeply inserted membrane-bound form is anchored in the membrane with majority of its residues interacting with the lipids. In addition to the “fusion active core” discussed above, we also observe direct lipid-interactions for other specific residues that might be targeted in future mutagenesis studies. Most of the hydrophobic residues, e.g., F817/I818, V826/T827/L828/A829, and F833/I834, are involved in binding and insertion into the membrane along with a few charged residues (Fig. 7B). Insertion of some of these residues is key to reducing hydrophobic exposure of the FP to the water environment, hence likely to increase the binding affinity of the FP. In addition, deeply inserted F817 and F833 are distinctive to and conserved among the SARS coronaviruses, which are currently the most infectious coronavirus species S1 . ^85^ Hence, we can speculate that F817 and F833 may function as an important factor in enhancing the infection rate of SARS-CoV2 by increasing the binding affinity of FP to the membrane. Future experimental studies will be necessary to further validate this hypothesis.

A recent complementary study has investigated the SARS-CoV2 FP membrane binding in the presence of Ca^2+^ ions.^86^ They observe different binding modes, including one in which the N-terminal helical segment binds the membrane along with L822/F823 penetrating the membrane, as well as a binding mode with insertion of F833/I834 into the lipid bilayer, which is in agreement with our findings. ^86^ Another interesting aspect is FP’s interactions with cholesterol. Although this study does not cover cholesterol specificity in binding of FP, binding affinity measurements for the FP in the presence of cholesterol supports that the FP favors interaction with cholesterol.^32^ Therefore, AS exemplified in Fig. 7A, we can speculate that the penetrating aromatic residues (F817, F823, F833, and Y837) in the membrane core are likely to interact with membrane cholesterol and increase the binding affinity.

In addition to the simulated peptide in this study (which we have referred to as the FP), there are other regions (named FP2 in SARS-CoV and FPPR in SARS-CoV2),^11,24–27^ which are implicated in membrane fusion. The fusion peptide proximal region (FPPR) of SARS-CoV2 downstream to the FP was later resolved and claimed to be involved in the structural rearrangement of the S-protein prior to membrane fusion.^27^ An internal disulfide bond within the FPPR, between C840 and C851, was observed and suggested to increase membrane-ordering activity.^25,27^ The membrane-ordering activity of the FP, due to the fusion active core, is significantly higher than the FPPR, and the activity of FP/FPPR together is only slightly increased compared to their activity separately. ^25^ Although our deeply inserted membrane-bound form supports the concept where both the FP and FPPR can interact with the membrane simultaneously as two subdomains,^25^ in order to characterize such a platform interacting with the membrane a longer peptide including the FPPR should be studied in future.

## CONCLUSION

COVID-19, which has emerged as a severe pandemic worldwide, calls for a need to accelerate the development of novel therapeutic intervention strategies. The S-protein of the SARS-CoV2 contains the key machinery necessary for the infection of human cell, including the FP, a highly conserved segment that inserts into the human cellular membrane and initiates the fusion of the virus. Yet, there are no post-fusion S-protein structures illustrating the binding of FP to human cellular membrane. In this study, using an extensive set of simulations, we describe how the SARS-CoV2 FP binds lipid bilayers representing mammalian cellular membranes and characterize, at an atomic level, lipid-protein interactions important for the stability of its bound state.

We capture different membrane-binding configurations from these simulations, which are classified using a detailed clustering analysis and based on geometrical evaluation of the peptide with respect to the membrane, resulting in three major membrane-bound configurations. Further analysis of these configurations in a mechanistic context taking into account the structural requirement of the entire S-protein, comparison of the results to previous experimental characterization of specific residues in the FP in the coronavirus family, and the degree of membrane engagement of the FP, all support the first binding mode (binding mode 1) to be the most likely and the most physiologically relevant form for the membrane-bound SARS-CoV2 FP.

Characterizing the mechanism of the fusion driving FP-host membrane interactions is key to our understanding of the critical steps involved in viral infection, paving way for potential development of novel therapeutics against SARS-CoV2, in addition to targeting the binding and interaction of the S-protein with the ACE2 host receptor. These include modulation of FP-membrane binding interface through small molecules with high affinity and specificity for this region of the S-protein, or inhibiting the key lipid-protein interactions observed.

Based on the suggested binding mode elucidated in our study, mutagenesis experiments can be designed to further confirm the role of the important residues implicated in membrane binding. Given the close similarity of the fusion peptides in coronaviruses in general, these results can also be applicable to infections caused by other members of this life-threatening family of pathogens.

## Author Contributions

DG, KK and ET designed the research. DG carried out all simulations, DG and ML analyzed the data. All authors wrote the article.

## Acknowledgments

This work was supported by the National Institutes of Health (Grants P41-GM104601 and R01-GM123455 to ET). MD simulations were performed using Blue Waters and computational resources provided by Microsoft Azure.

## Notes

### Competing Interest Statement

The authors have declared no competing interest.

### Summary of Updates

In this revised version of the manuscript, major changes include significant extension of all membrane-binding simulations from 100 ns to now 300 ns per replica, bringing the aggregate simulation time for the study to 9 microseconds. The clustering analysis has been accordingly updated and results in three major clusters for the membrane-bound form of the fusion peptide, with one representing the most biologically relevant form, which we discuss in detail and put in the context of previous studies. The Results/Discussion and main conclusions of the paper have also been substantially revised, accordingly.

