## Supplemental Figures for "Binding Mode of SARS-CoV2 Fusion Peptide to Human Cellular Membrane"

### Supporting Information

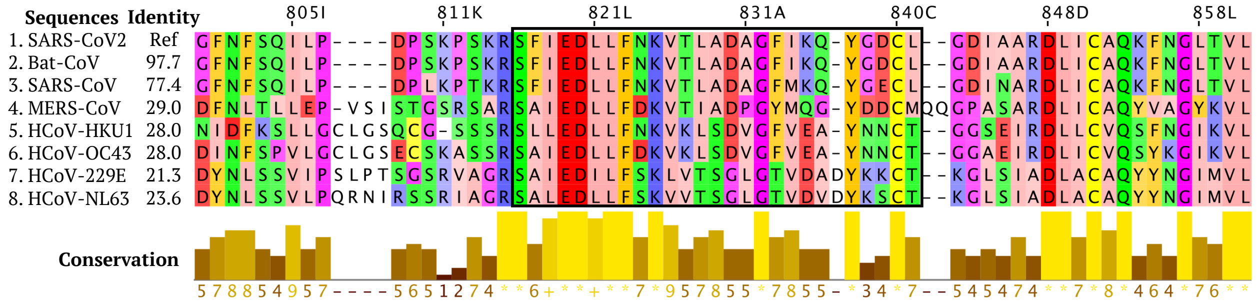


Figure S1: **Multiple sequence alignment of coronaviruses.** The S-protein sequence from SARS-CoV2 is aligned with the S-protein from closely related bat coronavirus RaTG13 (Bat-CoV) and six other human HCoV S-proteins. The residues are colored according to their physicochemical properties (Zappo coloring scheme), with aliphatic/hydrophobic residues (ILVAM) shown in pink, aromatic residues (FWY) in orange, positively charged residues (KRH) in blue, negatively charged residues (DE) in red, hydrophilic residues (STNQ) in green, conformationally special residues (PG) in purple, and cysteine (C) in yellow. The FP sequence used in this study is enclosed in a black box. The relative sequence identity of SARS-CoV2 S-protein with other sequences is shown (identity column). Conservation between the different sequences is visualized as histogram bars with their heights, and color variation from brown to yellow, reflecting the level of conservation of physicochemical properties in the alignment. Conserved and identical amino acid columns show the highest score of 11 and are indicated by '*' whereas conserved and similar amino acid columns show a score of 10 and are indicated by '+'. Other groupings are indicated by lower scores accordingly.

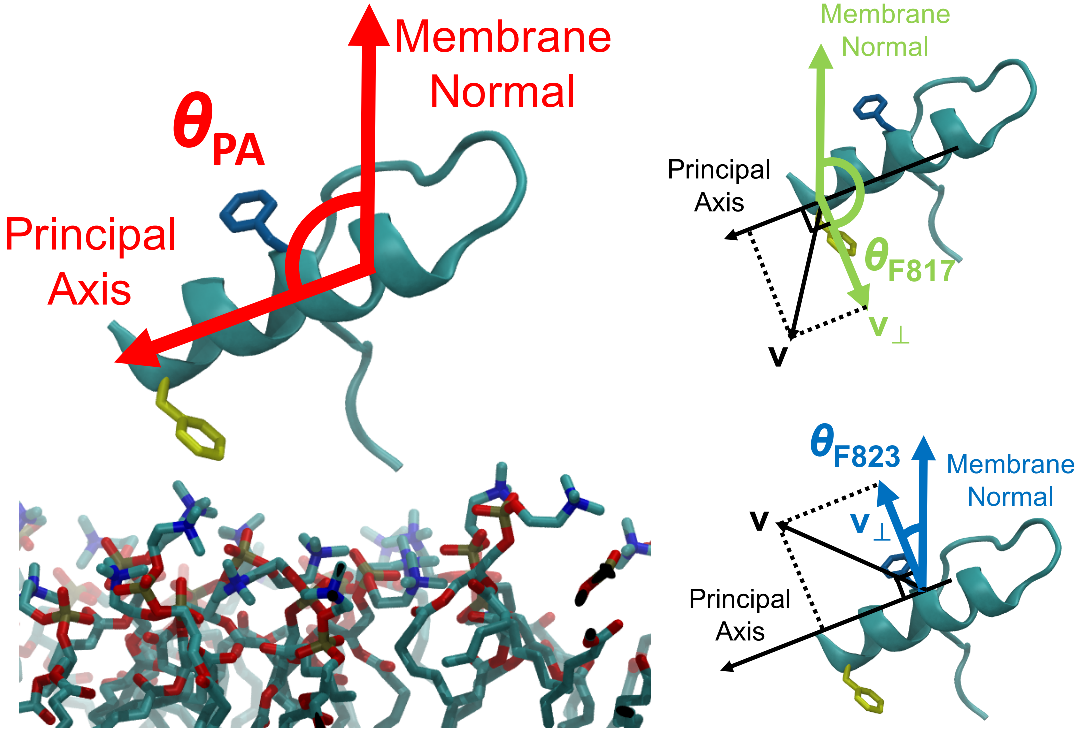


Figure S2: **Definition of the angles ($\theta_{\mathbf{PA}}$, $\theta_{\mathbf{F817}}$ and $\theta_{\mathbf{F823}}$) used to describe the orientation of the FP in its membrane-bound form.** The angle $\theta_{\mathrm{PA}}$ is defined as the angle between the membrane normal and the first principal axis (PA) of the helical segment (residue 816-823), which describes the tilting of the helical segment. Two auxiliary angles, $\theta_{\mathrm{F817}}$ and $\theta_{\mathrm{F823}}$, are identified to describe the orientation of two phenylalanine residues (F817 and F823) on the helix with respect to the membrane, which together with $\theta_{\mathrm{PA}}$ provide a more complete description of the orientation of the FP. These two angles are individually defined by the angle between the membrane normal and $\mathbf{v}_\perp$, the component of respective phenylalanine C$\alpha$-C$\beta$ vector (denoted as $\mathbf{v}$) perpendicular to the PA.

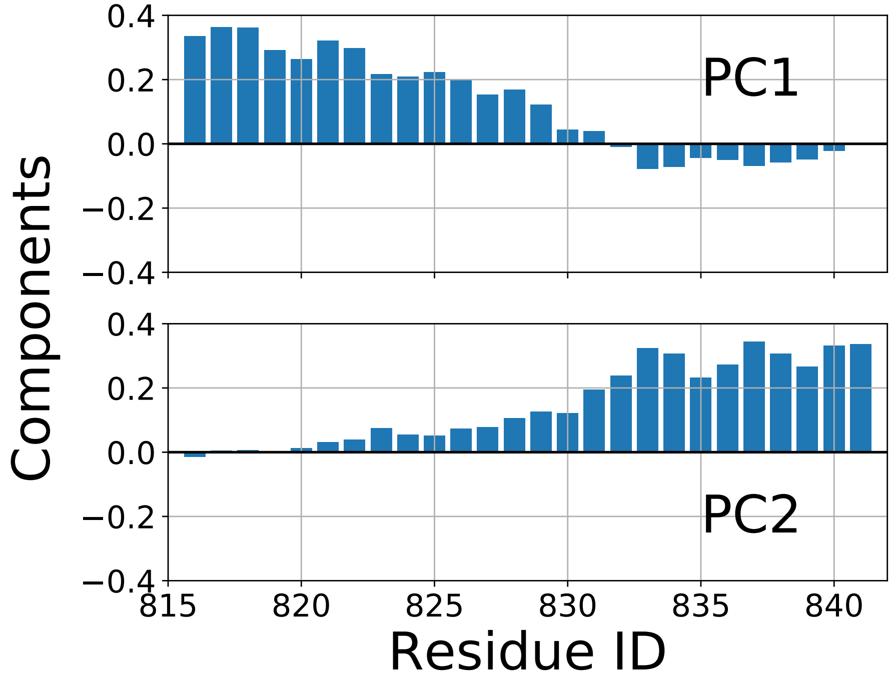


Figure S3: **The first two PCs defining the reduced dimension used to analyze the membrane-bound configurations of the FP observed in our simulations.** PC1 represents the level of membrane insertion of the $\alpha$-helical segment, whereas PC2 represents the level of membrane insertion of the C terminal loop segment.

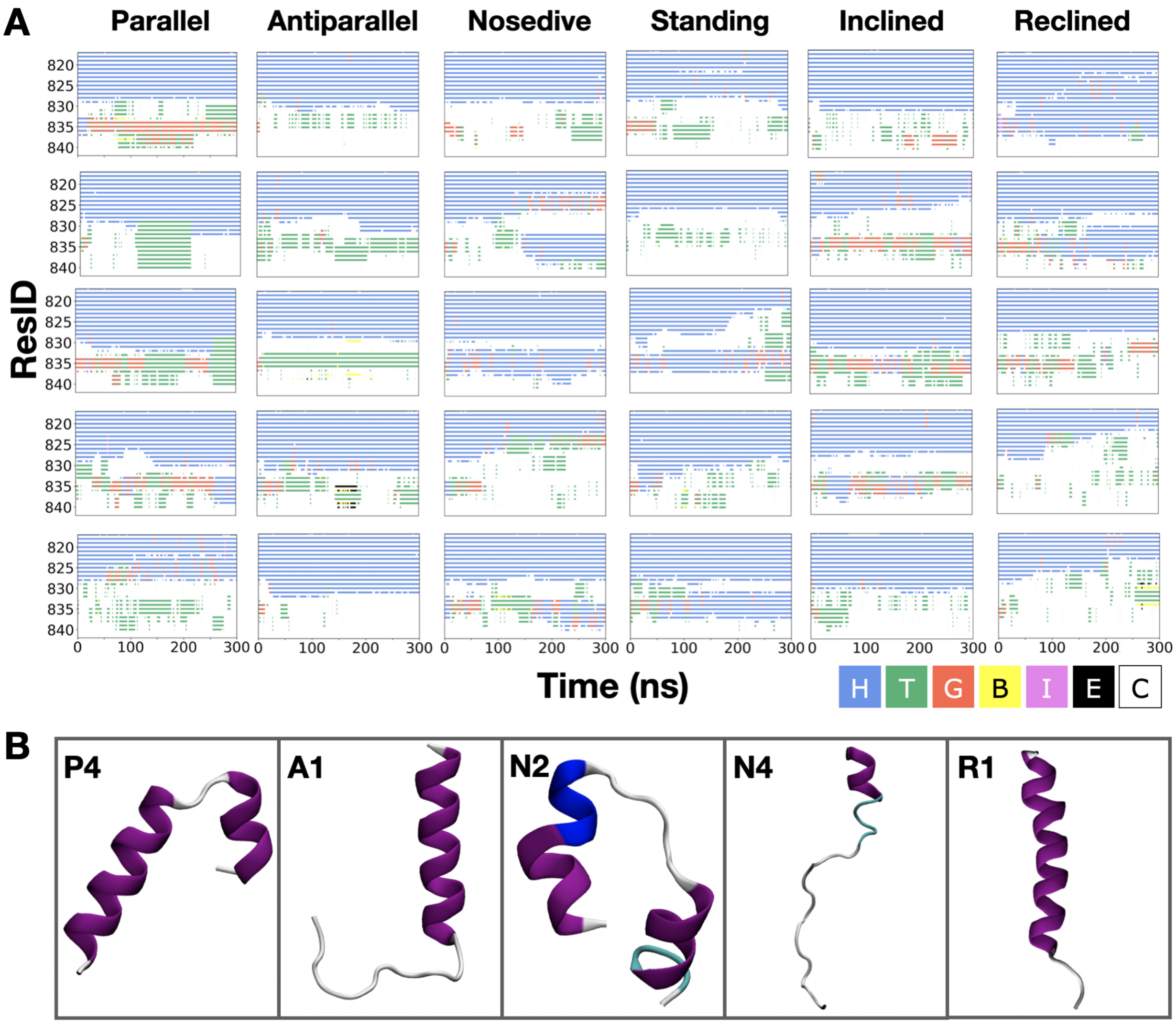


Figure S4: **Secondary structure analysis of the FP during membrane-binding simulations.** A) We monitored the evolution of the secondary structure in each replica to select the stable $\alpha$-helical part of the peptide. This allowed us to define the most stable segment of the structure to use for the definition of the orientational angles. The color keys for each type of secondary structure are H ($\alpha$-helix, blue), T (turn in $\beta$-sheets, green), G ($3_{10}$ helix, red), B (isolated $\beta$-bridge, yellow), I ($\pi$-helix, pink), E (extended strand in $\beta$-sheets, black), and C (random coil, white). B) Last-frame structures from some of the membrane-bound simulations representing the structural diversity of the FP. We observe hairpin structures in replicas P4 and N2, as well as unfolding (N4) and extension (R1) of initial helical structure. However, among the stably bound FPs, unfolding of the helical segment is uncommon.

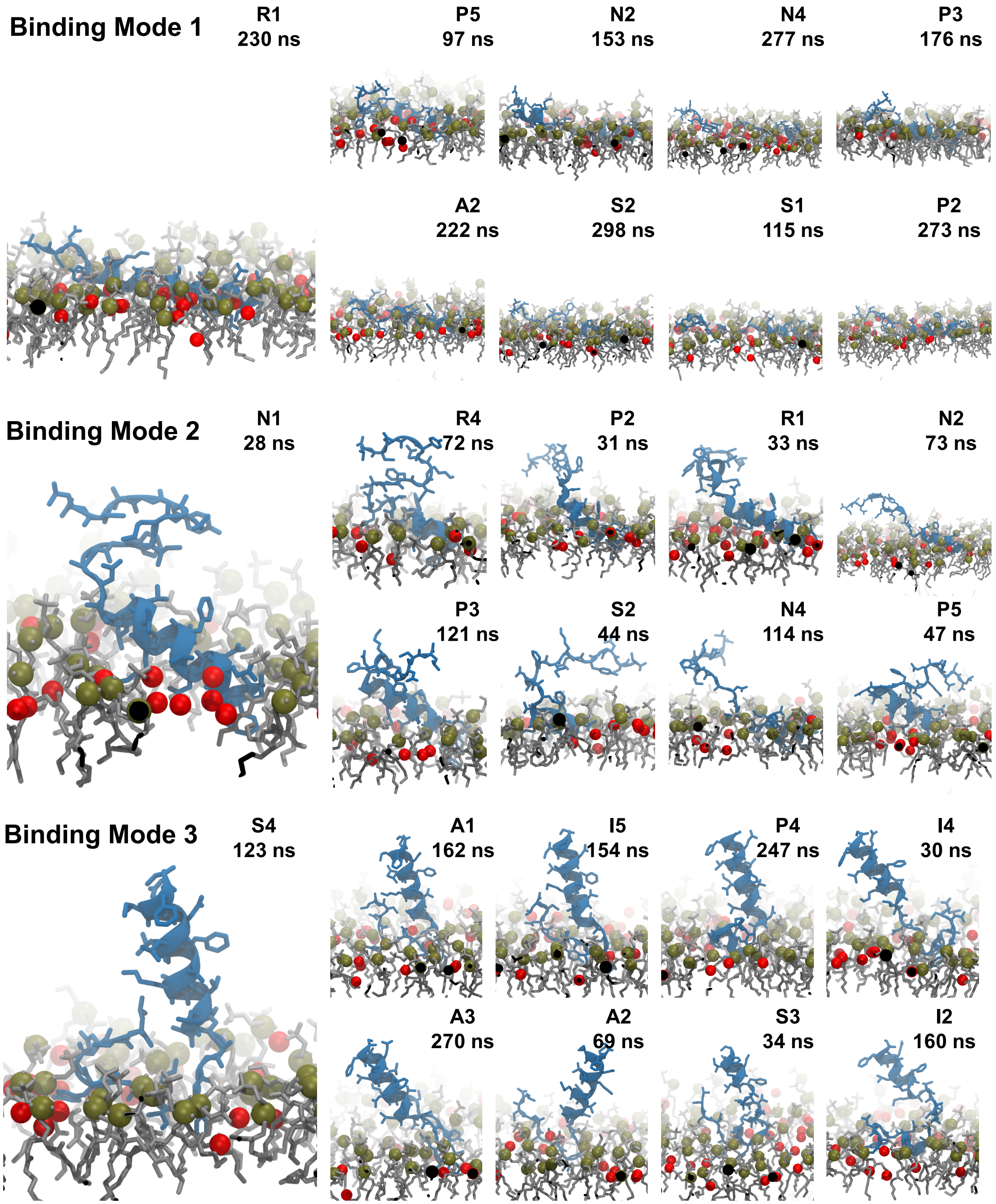


Figure S5: **Convergence of the three FP membrane binding modes.** The three representative structures (left), i.e., cluster centers, are shown with their closest eight snapshots (right) extracted from replicas of different independent simulation runs. FP side chains and PC lipids are shown as blue and brown licorice, respectively. Phosphorus atoms of PC lipids and oxygen atoms of cholesterol molecules are shown as brown and red spheres, respectively.
